# RTHybrid: a standardized and open-source real-time software model library for experimental neuroscience

**DOI:** 10.1101/426643

**Authors:** Rodrigo Amaducci, Manuel Reyes-Sanchez, Irene Elices, Francisco B. Rodriguez, Pablo Varona

## Abstract

Closed-loop technologies provide novel ways of online observation, control and bidirectional interaction with the nervous system, which help to study complex non-linear and partially observable neural dynamics. These protocols are often difficult to implement due to the temporal precision required when interacting with biological components, which in many cases can only be achieved using real-time technology. In this paper we introduce RTHybrid (www.github.com/GNB-UAM/RTHybrid), a free and open-source software that includes a neuron and synapse model library to build hybrid circuits with living neurons in a wide variety of experimental contexts. In an effort to encourage the standardization of real-time software technology in neuroscience research, we compared different open-source real-time operating system patches, RTAI, Xenomai 3 and Preempt-RT, according to their performance and usability. RTHybrid has been developed to run over Linux operating systems supporting both Xenomai 3 and Preempt-RT real-time patches, and thus allowing an easy implementation in any laboratory. We report a set of validation tests and latency benchmarks for the construction of hybrid circuits using this library. With this work we want to promote the dissemination of standardized, user-friendly and open-source software tools developed for open- and closed-loop experimental neuroscience.

## 1 INTRODUCTION

The study of neural systems dynamics is hindered by various factors. The first one is their intrinsic nonlinearity, since they process information in several interacting spatial and temporal scales and are affected by multiple transient adaptation and learning mechanisms. Also, from all magnitudes involved in these dynamics, just a few can be accessed simultaneously, making the system partially observable. The third factor is related to the use of the traditional stimulus-response paradigm in most experimental neuroscience research, which only allows to record the behaviour of the system under different stimuli and then to analyze the collected data offline. Thus, highly complex non-stationary neural activity, which has influence from the context and previous events’ feedback, can not be completely assessed. Closed-loop techniques provide an efficient way to overcome such difficulties by interacting online with the system, producing precise stimulus according to the recorded information and presenting valuable insights on transient neural processes. This paradigm allows more flexibility in the experiment and favours its automation, as well as the control of neural dynamics (Chamorro et al., 2012; Roth et al., 2014; Potter et al., 2014; Varona et al., 2016).

While closed-loop technology allows researchers to conduct online observation, control and interaction with neural elements, it also presents some drawbacks in its implementation. Some of these difficulties are related to the complexity of their experimental design, but also to the accomplishment of precise temporal restrictions, in the scale of milliseconds or lower, which often are required during data acquisition and stimulation in biological experiments (Christini et al., 1999; Muñiz et al., 2005, 2008, 2009). The capacity of a system to perform periodic tasks and respond to asynchronous external events in an strict time slot (neither sooner nor later) is known as real-time (Furht et al., 1991). Therefore, to ensure the compliance of the previously mentioned temporal margins in an experimental setup, real-time technology is needed. It is also important to mention that the term “real-time” may as well be used in neuroscience literature referred to online recording, feedback or control (Siegle et al., 2017). In this paper we always refer to strict temporal precision in the millisecond range.

Electronic components easily fulfill the speed and precision requirements needed for most real-time scenarios, so hardware-based implementations are one possible solution (Robinson and Kawai, 1993; Le Masson et al., 1995; Broccard et al., 2017). However hardware solutions are typically expensive or little programmable and manageable, especially if compared to software solutions, which offer maximum flexibility and a wide range of user-friendly frameworks. Moreover, nowadays personal computers have enough hardware capacity to comply with standard real-time restrictions. However, modern general purpose operating systems (GPOS), such as Windows, MacOS or GNU/Linux, are multitask environments with internal schedulers, which assign computer resources to different running tasks following specific policies (Stallings, 2012). These schedulers can not be controlled by the users, hence it can not be ensured that a given task will run without interruptions and, therefore, real-time can not be assured. In order to run protocols accomplishing a set of established temporal boundaries with software-based real-time, another system framework, known as real-time operating system (RTOS), is needed.

Numerous real-time tools for experimental neuroscience are already available. Some of them are hardware-based (Franke et al., 2012; Tessadori et al., 2012; Müller et al., 2013; Desai et al., 2017). Several software tools have been designed, particularly for dynamic-clamp electrophysiology experiments, both following soft-(Pinto et al., 2001; Nowotny et al., 2006; Linaro et al., 2014; Ciliberti and Kloosterman, 2017; Hazan and Ziv, 2017) and hard real-time (Christini et al., 1999; Dorval et al., 2001; Muñiz et al., 2005, 2009; Biró and Giugliano, 2015; Patel et al., 2017) approaches, using distinct platforms and RTOS, which have diverse purposes and architectures, hence presenting different advantages and disadvantages.

One example of closed-loop interactions can be found in hybrid circuits, which are networks built by connecting model neurons and synapses to living cells. They are a powerful tool to explore and characterize neural system dynamics, as well as a means to assess the role of specific circuit components, e.g. see (Yarom, 1991; Szücs et al., 2000; Pinto et al., 2000; Varona et al., 2001; Le Masson et al., 2002; Nowotny et al., 2003; Oprisan et al., 2004; Arsiero et al., 2007; Chamorro et al., 2009; Grashow et al., 2010; Brochini et al., 2011; Wang et al., 2012; Thounaojam et al., 2014; Hooper et al., 2015; Norman et al., 2016; Broccard et al., 2017). The most common paradigm to build hybrid circuits consists in using dynamic-clamp to read the membrane potential of a cell and, after computing a model using this voltage, inject the resulting current into the same or into a different cell (Robinson and Kawai, 1993; Sharp et al., 1993; Prinz et al., 2004; Destexhe and Bal, 2009; Nowotny and Varona, 2014). Neuron models range from simple mathematical approximations to more complex implementations based on Hodgkin-Huxley equations, which can reproduce biophysical behaviours with accuracy. Same thing happens for synapse models, which can cover from simple gap junctions using Ohm’s law to chemical connections defined by non-linear equations (Torres and Varona, 2012).

Beyond dynamical-clamp protocols, other stimulation and detection techniques that can improve their performance by using real-time technology are fMRI (Rana et al., 2016), optogenetics (Krook-Magnuson et al., 2013; Prsa et al., 2017), EEG setups (Arrouët et al., 2005; Sitaram et al., 2016), neuroprosthesess (Levi et al., 2018), or any kind of activity-dependent stimulation experiment, such as the ones that use simultaneous electrophysiological and video tracking (Muñiz et al., 2011), acute mechanical stimulation (Muñiz et al., 2008), electric signalling during behavior (Forlim et al., 2015; Lareo et al., 2016) or drug microinjection (Chamorro et al., 2009, 2012).

Due to the lack of flexibility in hardware real-time solutions, as well as RTOS heterogeneity and intrinsic difficulties in their use, many neuroscience researchers overlook this technology when designing closed-loop experiments. In this paper we introduce RTHybrid, a novel, multiplatform, real-time software neuron and synapse model library to build hybrid circuits. With this tool we aim to promote the use of standardized and user-friendly real-time software technology, available in different platforms, to favor the implementation of closed-loop experimentation in neuroscience research. We provide validation examples of this tool in the context of hybrid circuit implementations using dynamic-clamp, including detailed analysis and benchmarking of temporal precision in different RTOS.

## 2 MATERIALS AND METHODS

### 2.1 Real-time software

Real-time performance is often wrongly considered as a matter of speed, which is of course important, when it actually relies on temporal precision: actions must be delivered within a pre-established interval, neither sooner nor later. Many neurons follow precisely a similar behavioural pattern: slow activity (less than 1 kHz) but precise subcellular sequential dynamics or spiking coding.

When a computer performs a given task, there is always some latency between the moment when this task is expected to be accomplished and when it is actually done, as well as some jitter of these latency values, due to the performance of the operating system scheduler. Moreover, not all tasks are equally sensitive to high latency values or data loss, hence real-time software can be classified in two types: soft real-time, when some deadlines can be missed without performance degradation as long as some threshold is not exceeded (for example, an online music streaming service can lose some data packages and users will not notice it), and hard real-time, when all deadlines must be met or the system fails critically (computers controlling a nuclear plant or a satellite, for example) (Shin and Ramanathan, 1994). In this paper, the term real-time always refers to hard real-time.

Despite how differently a RTOS can be designed, the functioning of all of them rely on two elements: their scheduling algorithms, which are usually preemptive, meaning that they are able to interrupt a running process without its permission; and how they manage hardware interruptions (Abbott, 2006). Some of them are implemented from scratch, while others are based in existing GPOS’ kernels (Hambarde et al., 2014). Among the later, dual-kernel (a real-time microkernel is used along the standard one) and single-kernel solutions (the standard kernel is modified to support real-time) are common (Yaghmour, 2003; Dietrich and Walker, 2005).

There exist several commercial RTOS implementations, mainly designed for embedded systems, such as QNX Neutrino, VxWorks and Windows Compact Embedded, among others. Other proprietary real-time software solutions are MATLAB’s toolbox xPC and National Instruments’ LabVIEW Real-Time, which can achieve hard real-time precision if external hardware is used for the computation but otherwise are soft real-time, and Simulink Desktop Real-Time, which provides a hard real-time kernel (up to 1kHz sample rate) for executing Simulink models running Windows or Mac OS X. The main drawback of these tools is their high economic cost, which may be unaffordable for many research groups and laboratories. There are also open-source and free solutions to obtain an RTOS from Linux, an open-source GPOS, which avoid the stated inconveniences of commercial options and provide similar usability, or even better, in terms of performance (Aroca and Caurin, 2009; Hambarde et al., 2014). In this paper, three of these patches will be studied, comparing their performance and characteristics in relation to the task of building hybrid circuits with living and model neurons: RTAI, Xenomai and Preempt-RT.

RTAI (Real-Time Application Interface) (Mantegazza et al., 2000) was first developed in 1996, becoming one of the first and most widely used open-source real-time environments. It is based in a dual-kernel approach, using a real-time microkernel with a preemptive scheduler, which controls the interrupt requests (IRQ) and treats the standard Linux kernel as a low priority task. Another dual-kernel solution is Xenomai (Gerum, 2004), a project which was part of the RTAI project until 2005, when it become a fully independent tool. Other RTOS follow a single kernel approach, meaning that no auxiliary microkernel is used and that the standard one is modified instead to work in real-time. All changes made to Linux standard kernel for this purpose throughout the past decade are included in Preempt-RT (Dietrich and Walker, 2005).

### 2.2 Experimental setup

#### 2.2.1 Benchmarking tests

Performance on the three RTOS described in the previous section, RTAI, Xenomai and Preempt-RT, was measured and compared among them and also to a GPOS with no real-time capabilities. Specifications of the computers used for these tests can be found in Table 1. Most modern computers currently have multicore processors i.e. one component with several independent processing units. Linux also allows to isolate specific cores, so the scheduler will not assign them any task, and to manually bound an specific task to this empty core. We have also analyzed how this core isolation affects the performance in both real-time and non real-time implementations.

**Table 1.** Software and hardware specifications of the computers used on the benchmarking tests for each operating system and RTOS used in this study. Note that the computer used for non-real-time, Preempt-RT and Xenomai 3 tests was the same (we will call it Computer 1 hereafter) and the one for RTAI was different (Computer 2).

|  | Operating system | Kernel version | RAM | Processor | Cores |
| --- | --- | --- | --- | --- | --- |
| **Non real-time** | Debian 9 | 4.9.0-4 | 16 GB | Intel Core i7-4790 3.6 GHz | 4 |
| **Preempt-RT** | Debian 9 | 4.9.0-4 | 16 GB | Intel Core i7-4790 3.6 GHz | 4 |
| **Xenomai 3.0.5** | Ubuntu 16.04 | 4.9.90 | 16 GB | Intel Core i7-4790 3.6 GHz | 4 |
| **RTAI 3.4** | Ubuntu 10.04 | 2.6.34.5 | 4 GB | Intel Core i7-2600 3.4 GHz | 4 |

The benchmarking procedure consisted in a latency test, measuring the time difference between when an action was expected and when it really happened (see Fig. 1). The benchmarking program^1^ consisted in a periodic loop with a frequency of 20kHz that sent a digital 0 or 1 to a digital acquisition (DAQ) device alternately on each iteration, thus producing a 100*μs* square-wave signal. After sending the corresponding value the program slept until the next interval arrived: the time interval between the real and the expected awaking time was the measured latency. An *Agilent MSO7104A* oscilloscope was used as an external temporal reference and *stress*^2^, a workload generator software, was utilized to create a worst case scenario, with all processor cores and the file Input/Output system running at full capacity. To send the signal to the oscilloscope a National Instruments PCI-6251 board and a BNC-2090A DAQ device were used.

**Figure 1.**
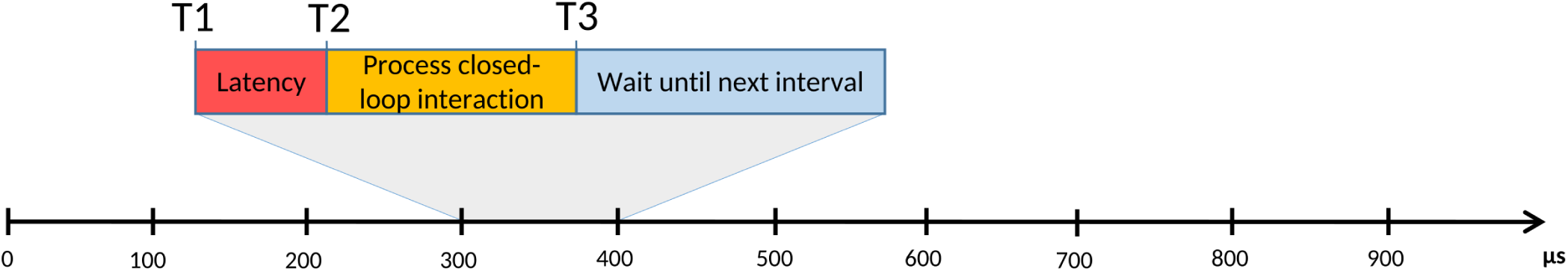
Representation of a typical real-time process, consisting on a succession of fixed and equal duration time intervals that are repeated on each iteration of a periodic loop. For any of these intervals, T1 is the time when it should start, but, due to the system capacity or to its resource management algorithms, the beginning can be delayed until T2. The difference between the real start time (T2) and the expected one (T1) is known as latency. After T2, the process computes its iterative task, which finishes at T3. If T3 happens before the next interval start time, the process waits until that moment. On the other hand, if T3 occurs later than the expected start time of the next interval, as a result of a high latency value, a long computational time or both, it would cause a failure on the real-time system. Real-time tasks can be classified according to their tolerance of such events: soft real-time means that some deadlines can be missed under a certain limit, while hard real-time does not tolerate any failure.

#### 2.2.2 RTHybrid validation tests

In order to validate the proper functioning of RTHybrid, it has been tested in a real experimental environment. Hybrid circuits were built by connecting bidirectionally a neuron model simulated by the software and a living neuron, using chemical graded model synapses. For these tests, a computer with a 4-core Intel Core i7-6700 3.40 GHz processor and 16 GB RAM memory, as well as a National Instruments PCI-6251 board with a NIBNC-2090A DAQ device, were used. Measures of the time taken by each task in the interaction cycle were also conducted to evaluate their contribution to minimum and maximum computational cost for each of the real-time platforms analyzed.

The membrane potential of the biological component of the experiment was recorded using *in vitro* electrophysiology in neurons of the pyloric central pattern generator (CPG) of adult *Carcinus maenas*, bought in a local fish store and kept in artificial sea water. Before the dissection, the crab was anesthetized by introducing it in the freezer for 20/30 minutes. The stomatogastric ganglion, dissected following the standard procedure, was attached to a Petri with Sylgard cold saline dissolution (13-15°C kept by a microcontroller and always perfused) using pins. The saline had the following composition (in mmol/l): NaCl 433, KCl 12, CaCl_2_.2H_2_O 12, MgCl_2_.6H_2_O 20, Hepes 10. To adjust the pH to 7.4-7.6, NaOH 0.1 M was added. Neurons were identified after desheathing the ganglion by their membrane potential waveforms and their corresponding spike times in nerves. Intracellular recordings were performed using 3 M KCl filled microelectrodes (50 *M*Ω) and a DC amplifier (ELC-03M, NPI Electronic, Hauptstrasse, Tamm, Germany). For details on the preparation see (Elices et al., 2018).

## 3 RESULTS

### 3.1 Real-time benchmarking and comparison

#### 3.1.1 Computers internal clock validation

Tests were conducted to certify that the internal clocks of the systems specified in Table 1 were capable of working with the required microsecond precision. These consisted in three 10-second tests on each platform, generating a square-wave signal of period 100*μs*, which was recorded with an external oscilloscope. When compared to the signal period measured by the computers, the oscilloscope registered a ±10*μs* inaccuracy in the precision of the computers’ clocks (see Table 2). This is an acceptable margin for our purposes with sampling rates for the hybrid circuit electrophysiology ranging from 10-20 kHz.

**Table 2.** Results of the internal clock precision tests for the computers described in Table 1, in microseconds. A 100*μs* signal, generated by the computers, was recorded with each computer and an external oscilloscope. Minimum, maximum, mean and standard deviation values of the recorded signal period are displayed at the table. When compared to the signal period measured by the computers, the oscilloscope registered a ±10*μs* inaccuracy in the precision of the computers clocks.

|  | Computer 1 |  |  | Computer 2 |  |  |
| --- | --- | --- | --- | --- | --- | --- |
| μs | Min | Max | Mean ± Std | Min | Max | Mean ± Std |
| Measured by computer | 99.5 | 100.5 | 100.0 ± 0.1 | 99.9 | 100.1 | 100 ± 0 |
| Measured by oscilloscope | 90.0 | 110.0 | 100 ± 3 | 90.0 | 110.0 | 100 ± 3 |

#### 3.1.2 RTOS usability comparison

RTAI, Xenomai and Preempt-RT were studied and compared in terms of installation requirements, usability and user-friendliness. This analysis is summarized in Table 3.

##### RTAI

Installation requires to patch a vanilla Linux kernel with the RTAI patch and then compile it, which is a long and tricky operation, even for experimented users. The last version of the official installation guide dates from 2008 (Monteiro, 2008). Utilization of the real-time functions provided by the platform is done through its own API, which is very powerful and complete, but documentation and examples are scarce. The safest way to achieve real-time is implementing the programs as kernel space modules, which carries many impediments. User space real-time can be reached using the LXRT library, although its use is discouraged for non-senior RTAI programmers (Racciu and Mantegazza, 2006). Currently, this tool is still maintained, but not regularly: version 4 last maintenance was in 2013 and in May 2017 version 5 was released, followed by the 5.1 patch in February 2018.

##### Xenomai

Patching and compiling a Linux vanilla kernel is the only way of installing Xenomai and its developers provide an up-to-date guide. Since Xenomai’s main purpose is to offer an open-source alternative to proprietary RTOS, it includes different APIs, intended to emulate other environments and libraries, such as VxWorks, pSOS+ and even POSIX. All of them, as well as their own API, are accessible from user space. Complete and updated documentation is available for all APIs. Although the user community is not very large and there are not many examples to be found, there is a mailing-list where questions can be asked. The project is currently active (Xenomai 3 was released in October 2015) and it is maintained and updated frequently. An Ubuntu 16.04 distribution already patched with Xenomai 3.0.5 can be downloaded from our website^3^.

##### Preempt-RT

Similarly to the dual-kernel implementations, the typical way to install Preempt-RT is by patching and compiling a vanilla kernel, following the instructions that can be found at the official website, these being are significantly simpler than the ones for RTAI and Xenomai. An alternative possibility for Debian distributions is to install it from its repositories as any other package. Despite the real-time patch, the system is still a normal Linux, so the standard POSIX library can be used and all its documentation is valid. In 2015 the project was transferred to The Linux Foundation, becoming the “official” Linux real-time solution.

**Table 3.**
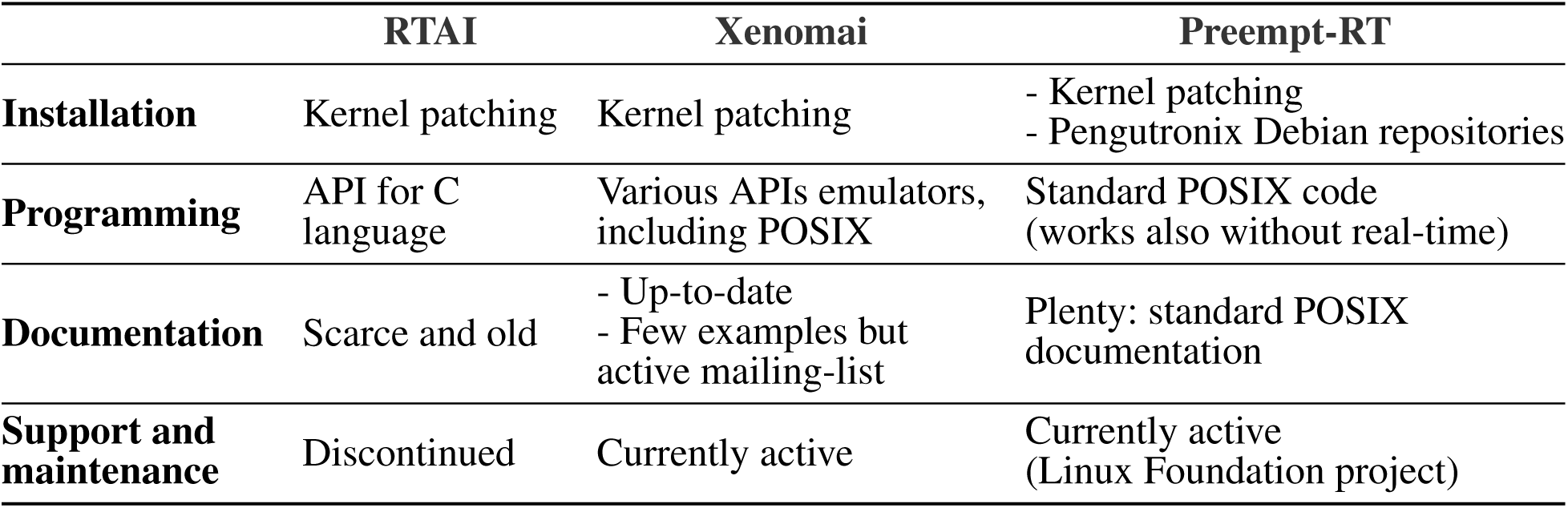
Usability and accessibility characteristics for each real-time solution explored in this study.

#### 3.1.3 RTOS benchmark analysis

RTAI, Xenomai and Preempt-RT were tested using the method described in 2.2.1. Each trial consisted in a five minute run of the test program under stress, with a frequency of 20kHz, and was repeated twice on every platform. We measured the maximum, minimum and mean latency values, as well as the jitter, and the results can be seen in Fig. 2. Distribution of latency values during these tests is shown in Fig. 3.

RTAI obtained the best performance scores, getting 3.65as maximum latency, even running on an older machine. Xenomai was not far from this performance, with a maximum latency of 5.66*μs*. Preempt-RT had slightly worse results, reaching a maximum latency of 15.94*μs*, but still acceptable for our purposes. The system without real-time is not reliable when millisecond precision or below is required, as it goes over the millisecond barrier in these tests (indicated in red in Fig. 3). Nevertheless, as mentioned in section 2.2.1, Linux operating systems allow to isolate a processing core and bind a specific task to it. In this scenario, the task running over the isolated core will never be interrupted by any other users tasks, but system processes can still use this core. When this is done in RTAI or Xenomai it has little impact on their already good latency values, but we observe a remarkable improvement of latency values with both non real-time and Preempt-RT operating systems. Without an RTOS, it can be an useful tool in soft real-time environments.

**Figure 2.**
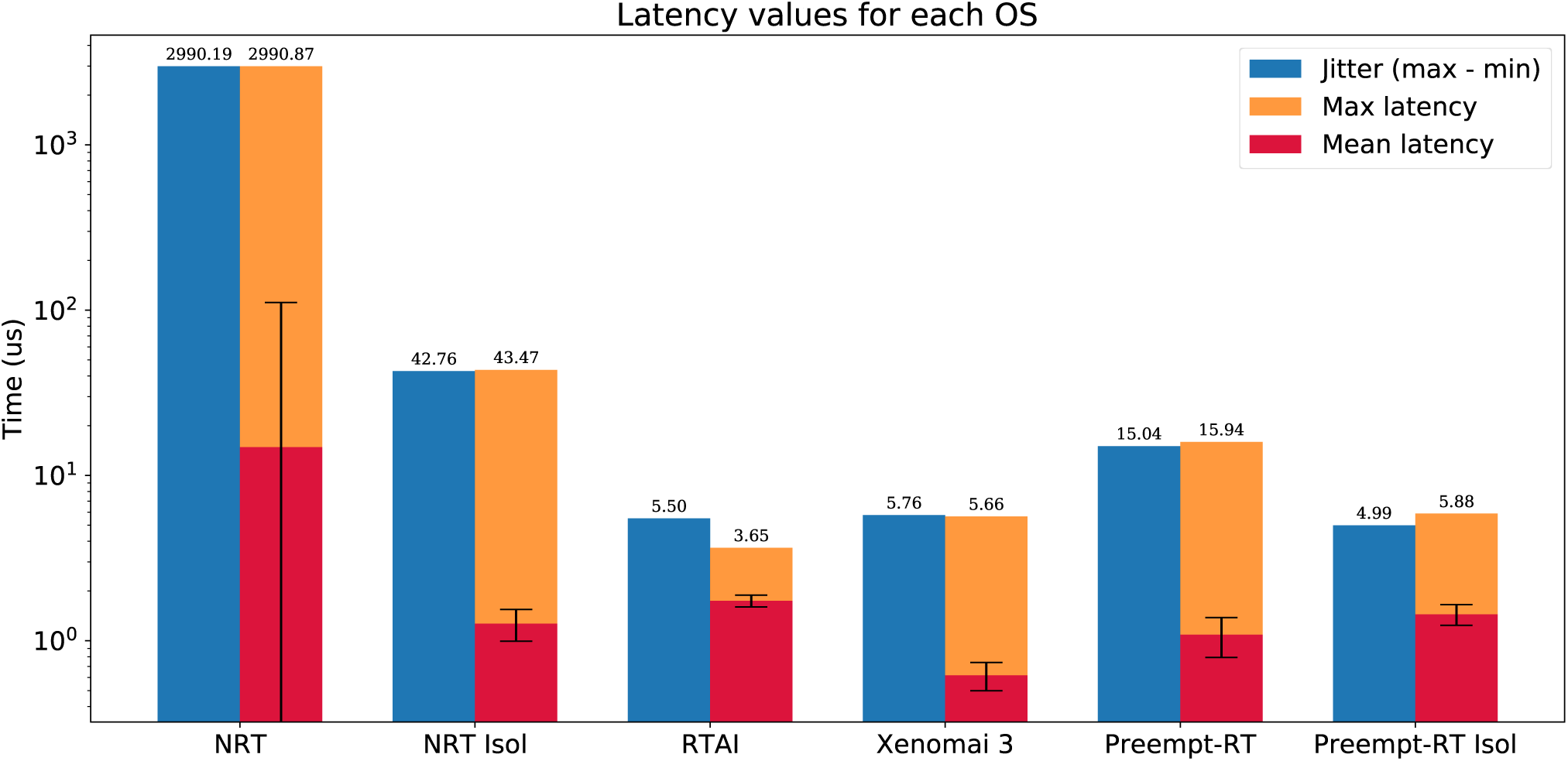
Summarized results of the real-time benchmarking tests for each OS: No real-time (NRT), No real-time with an isolated core (NRT Isol), RTAI, Xenomai 3, Preempt-RT and Preempt-RT with an isolated core (Preempt-RT Isol). Time axis is represented in log-scale. Blue bar represents the jitter, calculated as the difference between the maximum and minimum latency. Orange bar represents the maximum latency. Red bar is the mean latency, error bars indicate the standard deviation. Numbers on top of each bar correspond to the largest jitter and latency, respectively. For 20kHz trials the real-time constrain is 50*μs*, which is only exceeded in this case by the OS with neither real-time capabilities nor an isolated core.

Unfortunately, regarding the analyzed RTOS, the better their performance is, the worse their usability and user-friendliness. As summarized in Table 3, RTAI is quite difficult to install and use, even for experimented users, while Preempt-RT is the most accessible, since there are not many differences with a normal GPOS. Due to this results, RTHybrid was developed to run over both Xenomai and Preempt-RT to balance performance and user-friendliness.

### 3.2 RTHybrid

#### 3.2.1 Software design and implementation

RTHybrid is designed to be an user-friendly and accessible real-time tool for any researcher, regardless of their budget or computer science and programming knowledge. It is an open-source project that can be downloaded for free from its Github repository^4^. Any Linux distribution is supported, including those running Xenomai 3 and Preempt-RT real-time patches. The code is written using C/C language and compiled with GCC 6.3 and QMake 3.0. Relevant information regarding the hybrid circuit experiment, such as neuron (both living and model) and synapse types employed, parameters and latency values is registered in log files. Detailed instructions on how to download, configure and install both RTHybrid and all its dependencies are provided at its user manual^5^.

**Figure 3.**
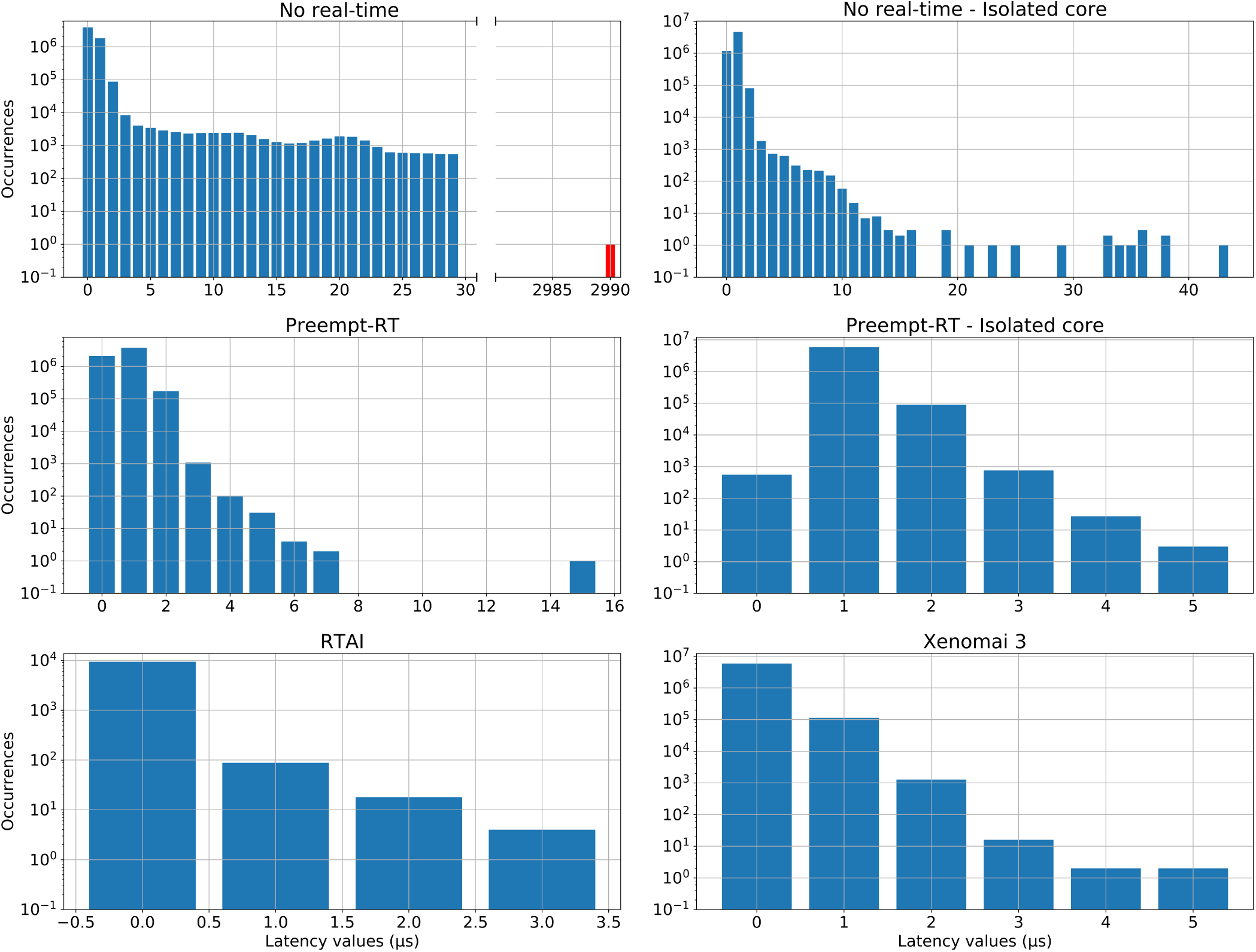
Distribution of latency values for each operating system during the real-time benchmarking tests. Red bars correspond to latency values that have exceeded the 50*μs* limit established for 20kHz trials, which is the case for the “No real-time” scenario. Vertical axes are in log-scale. This analysis was performed on the same data than Fig. 2.

Three processing threads have been used to build the program’s architecture in order to address an optimized real-time software implementation (see Fig. 4). Two execution modes are available: graphical user interface (GUI) mode or script mode. In the former, an intuitive GUI (see Fig. 5) is displayed when the application is launched, where the user can select the desired models and modify all their parameters as well as set the experiment configuration (duration, DAQ channels, sampling rate, etc). This GUI has been designed using the Qt 5.10 framework^6^. The second option allows to automatically load all experimental protocols and parameters from an XML text file^7^, without using the GUI. In this case, various experimental protocols can be automatically executed one after the other using scripts. Whatever option is chosen, the first and main process is in charge of gathering the information from either source and starting two new threads with it.

**Figure 4.**
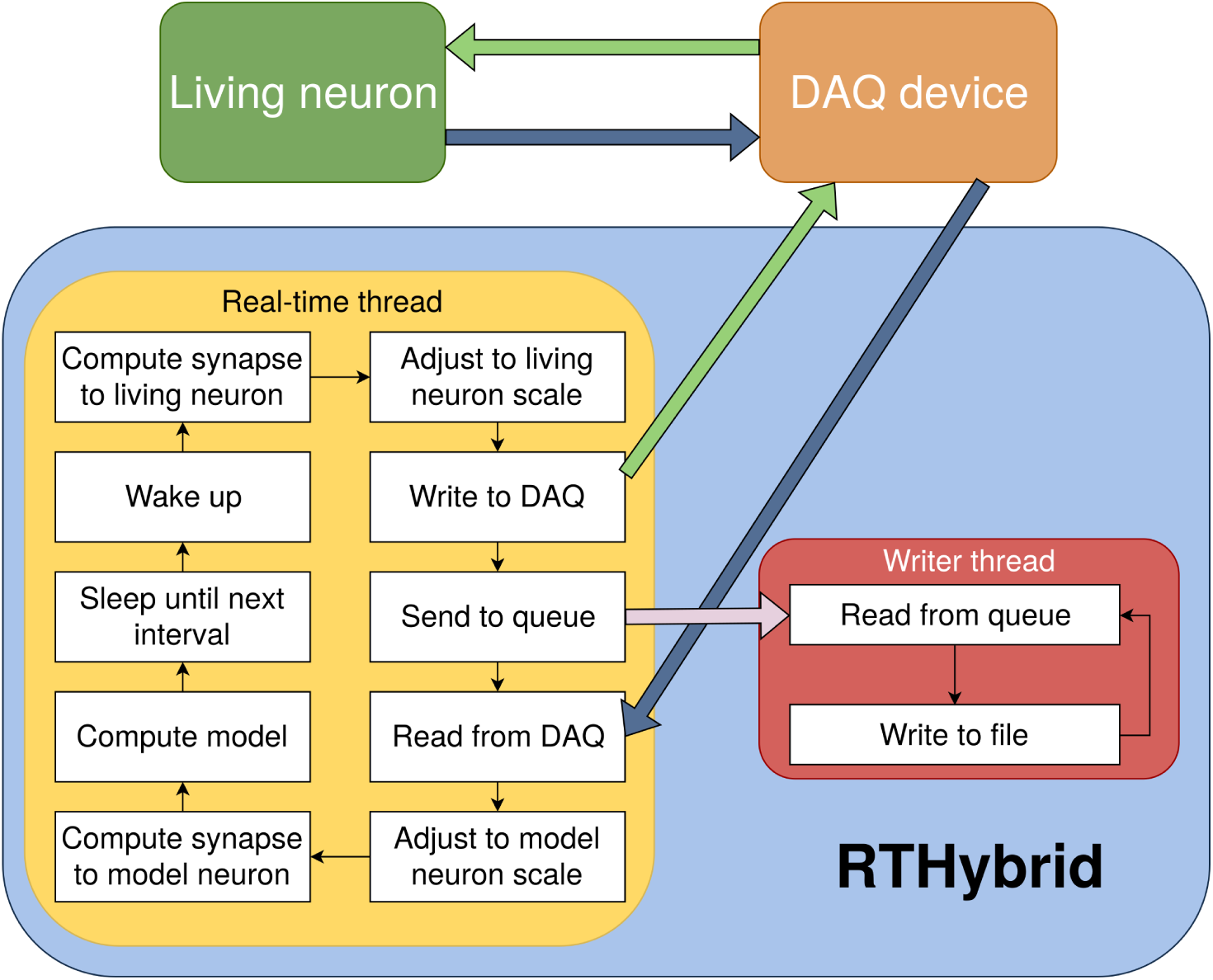
Diagram of RTHybrid architecture. The computer receives the membrane voltage from the living neurons through the DAQ device, computes the neurons and synapses models and sends current back to the biological cells. The main program process creates two sub-threads, *rt_thread* and *writer _thread*, to manage the different tasks of the program. Both threads communicate through inter-process communication (IPC) message queues.

The second thread performs all tasks that need to be completed with real-time precision in a periodic loop. Within each interval of this loop, key operations are fulfilled: interaction with the DAQ device, synapse and neuron models computation, automatic calibration, drift adaptation, etc. (see Fig. 6). Communication with DAQ devices is achieved through Comedi open-source drivers for National Instruments’ and several other manufacturers’ hardware (or Analogy drivers in the case of Xenomai) (Schleef et al., 2012). Automatic calibration and experiment automation algorithms are included to deal with the differences between models and living neurons in terms of temporal scale and amplitude (Reyes-Sanchez et al., 2017). They also cover other possible experimental complications such as the presence of signal drift.

Some tasks are computationally too expensive to be carried out within the temporal restrictions established in a real-time interval, including essential ones as writing data to a file or printing it on screen. A third thread is used to write experiment data to files without disturbing the real-time performance. An inter-process message queue is utilized to send information from one thread to the other. Since it is not a real-time process, it will wait until there are enough computer resources available, and only then reads from the queue and stores the data into the files.

**Figure 5.**
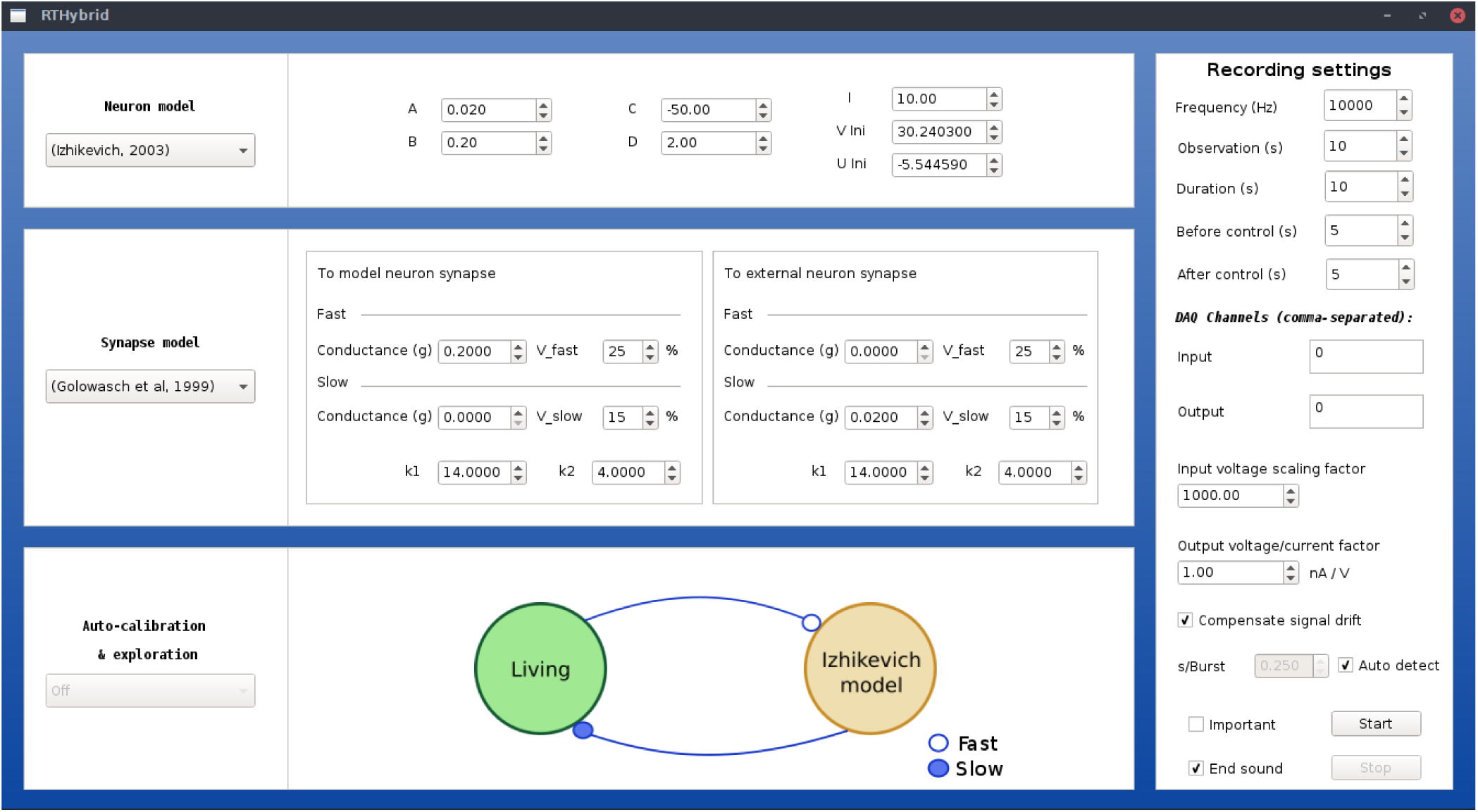
Illustration of RTHybrid graphical user interface to build hybrid circuit interactions. Users can select neuron and synapse models, as well as their parameters, and the experiment settings, such as input/output DAQ channels, sampling frequency, duration, etc.

#### 3.2.2 Model library

RTHybrid includes a customizable library of neuron and synapse models to build a wide variety of hybrid circuits. Table 4 shows the currently included neuron models, which have been selected due to their rich intrinsic dynamics and suitability to build hybrid circuits. All of them have different characteristics and require also specific adaptations to work in real-time, which are handled automatically by auto-calibration algorithms (Reyes-Sanchez et al., 2017). Several numerical integration methods can be chosen by the user, including Euler, Heun, Order 4 Runge-Kutta (Press et al., 1988) and (6)5 Runge-Kutta (Hull et al., 1972). Table 5 lists the types of synapse models included in the library at the moment and the different input parameters that they require to work in a hybrid circuit implementation.

**Table 4.**
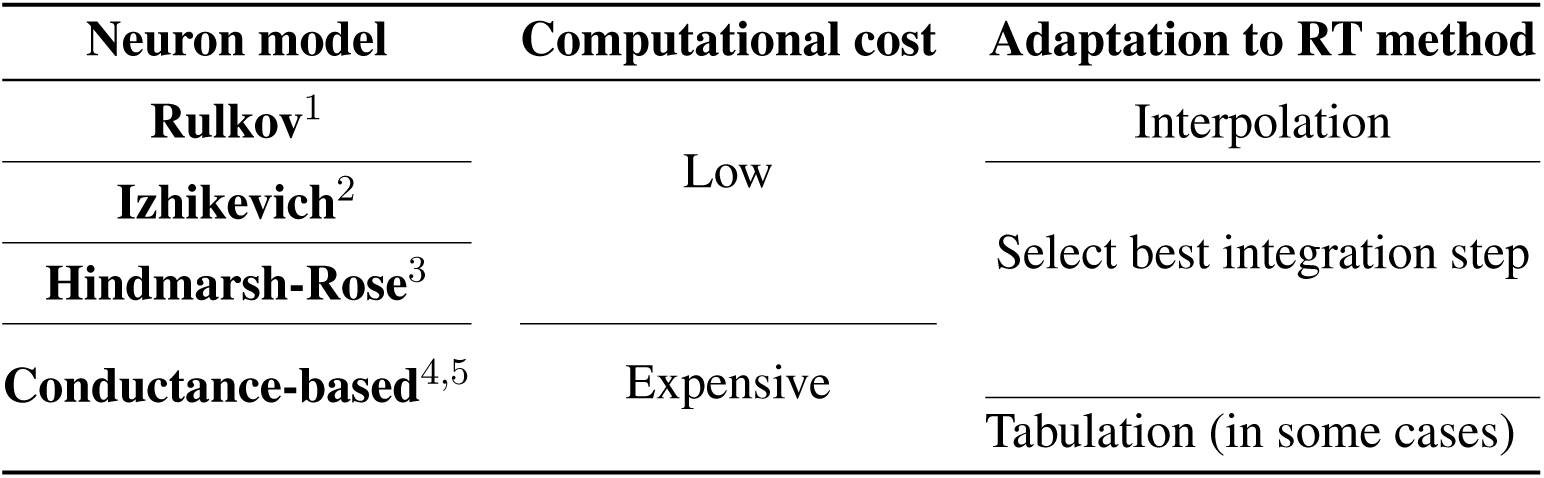
Currently available neuron models in RTHybrid. Different models have different computational costs and need to be adapted to real-time performance using distinct methods. ^1^Rulkov, 2002) ^2^(Izhikevich, 2003) ^3^(Hindmarsh and Rose, 1984) ^4^(Ghigliazza and Holmes, 2004) ^5^(Wang, 1993).

**Table 5.**
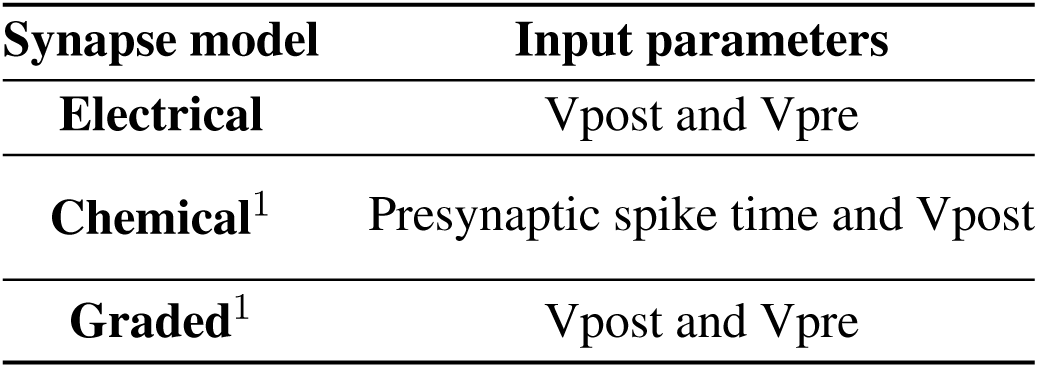
Currently available synapse models in RTHybrid. Different kinds of model may require different input parameters to work in a hybrid circuit configuration. V refers to membrane potential of a living or model neuron. ^1^(Golowasch et al., 1999).

Models described with differential equations require minimum integration steps and specific adaptations to match the duration of events such as spikes or bursts in a given living neuron. This is the case of models such as (Izhikevich, 2003; Hindmarsh and Rose, 1984; Ghigliazza and Holmes, 2004; Wang, 1993). Other neuron models, such as the Rulkov map (Rulkov, 2002), generate activity events such as bursts using a little number of points. In this case, interpolation of the values produced by the model is required to match the living neuron’s event resolution at the chosen sampling rate. Synapse models also present differences in their implementation, both in the input parameters required and the computational cost. A simple linear electrical synapse model is included along with a more complex chemical graded synapse one (Golowasch et al., 1999). New models can be easily added to the library using C language.

Figure 6 illustrates the average time consumed by each task at each iteration of the loop executed by RTHybrid real-time thread. This study was performed for several models of the RTHybrid library. The models were run at a 10kHz frequency for five minutes (300 seconds), i.e., each model test contained 3 million intervals of 100*μs* duration. Neuron models were bidirectionally connected through a chemical synapse model to a hardware-implemented Hindmarsh-Rose model that generated bursting activity at the same characteristic rate of a pyloric CPG cell (Pinto et al., 2000). Synaptic conductances were set to *g* = 0.02*μS* and all other parameters from neuron and synapse models were fixed to produce bursting behaviour. Burst duration for the models was set at one second per burst. The computer used for these tests was the one referred as Computer 1 in Table 1, with both Preempt-RT and Xenomai 3 and the same DAQ device and board described in section 2.2.2. Models selected for this analysis included one low-cost differential equation implementation (Hindmarsh and Rose, 1984), a fast map model (Rulkov, 2002) and two conductance-based models with exponential non-linearities (Ghigliazza and Holmes, 2004; Wang, 1993).

Variability of the time consumed by the operations performed inside each interval (see Table 6) is constrained by the use of RTOS, being Xenomai 3 more effective in this task than Preempt-RT. Xenomai 3 achieve much lower latency values, but the use of the Analogy framework, instead of Comedi, translates into a higher cost when interacting with the DAQ device. Some models have on average low computation times which may increase occasionally due to their nonlinearities or to a non stable regime for the chosen integration method. This is the case for the Hodgkin-Huxley type model from (Wang, 1993), which on less than ten occasions on every trial, out of the three million iterations described previously for Fig. 6, had an abnormally high computation time. Other models can always be computationally expensive because of different reasons such as their high dimensionality or multi-compartmental nature. One way to reduce their integration time and make them more efficient and suitable for real-time environments is to tabulate their nonlinearities. When the operations within an interval exceed the temporal margin established, RTHybrid tries to minimize the negative impact on the system by taking time from the next interval and reducing the sleeping period, as can be seen in Fig. 7. The presence of latency values above the real-time threshold can be detected from the log file.

**Table 6.**
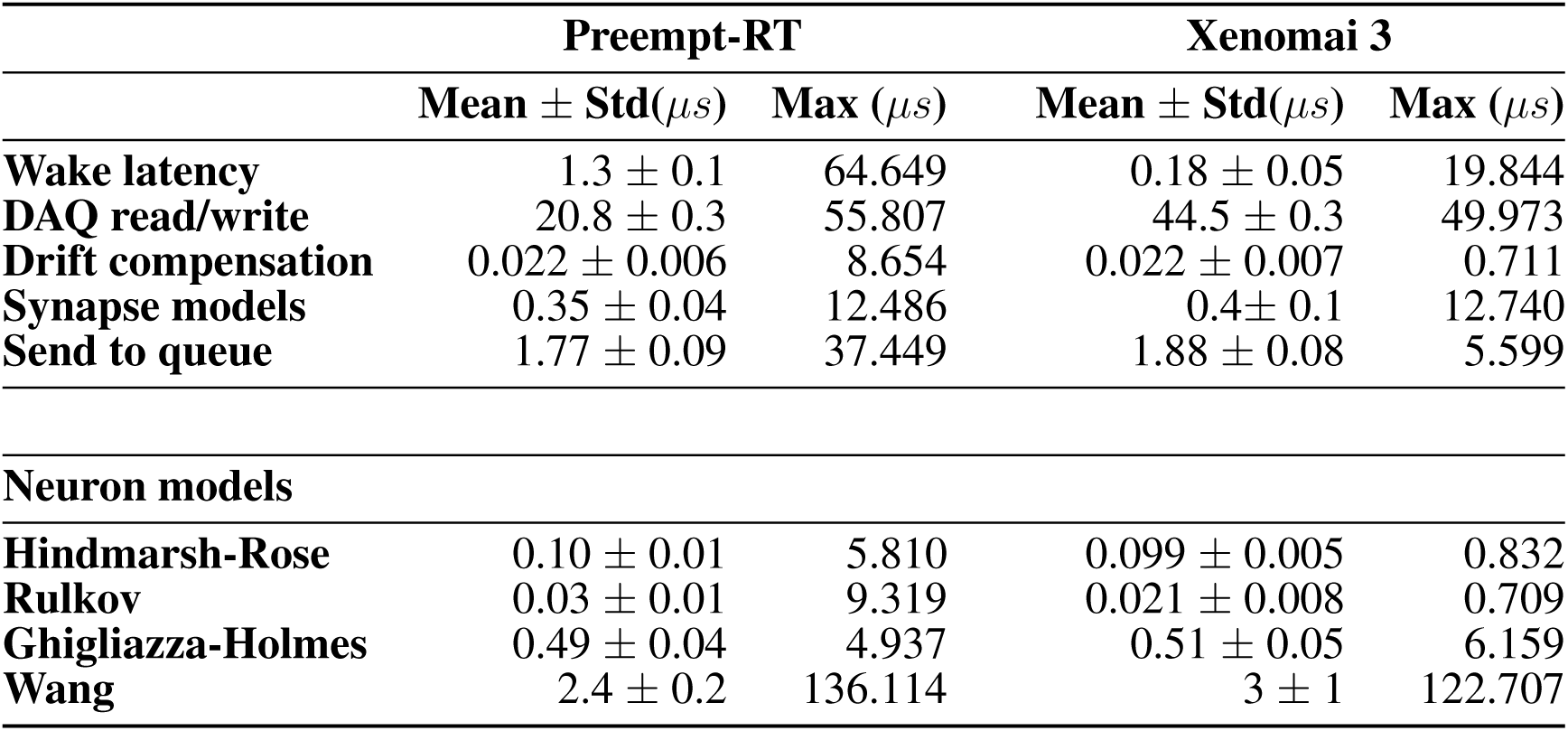
Duration analysis for each operation performed inside the real-time cycle described in Fig. 6 on every iteration. Mean and maximum times are shown in microseconds. The use of RTOS constrains the variability of these times, with Xenomai 3 being more efficient at this than Preempt-RT. Duration of each model computation differs greatly from the others due to their distinct mathematical descriptions.

**Figure 6.**
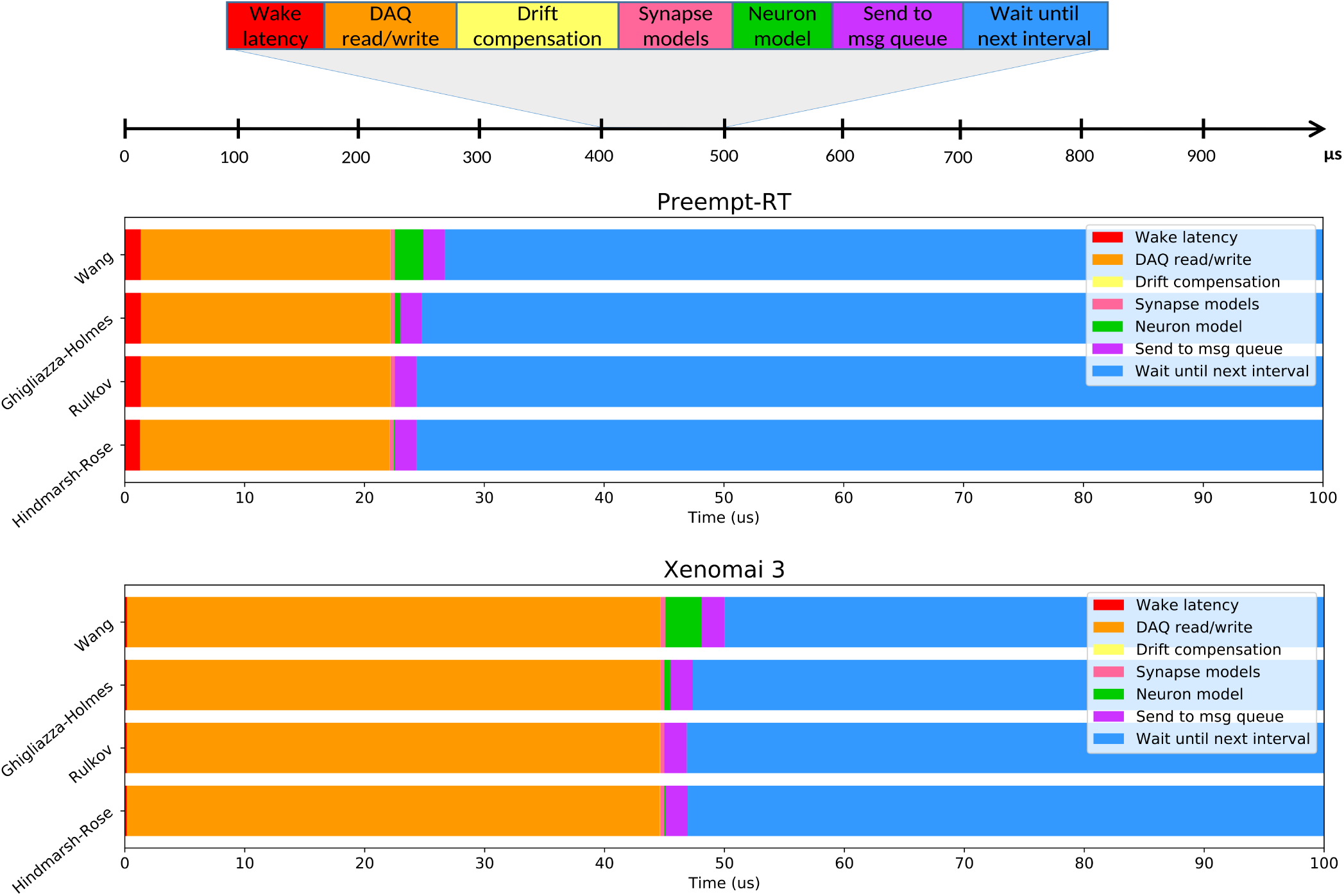
Average time usage for each operation over RTHybrid real-time intervals when different neuron models are computed. Top panel illustrates the operations that are performed inside each iteration of the loop executed at the real-time thread of RTHybrid: the process wakes up, interacts with the DAQ device, performs (if activated) the drift compensation, computes the synapse models, calculates a new point (or points) of the neuron model, sends the message with data for the writer thread to the queue and sleeps until the expected beginning of the next interval. Middle and bottom panels show the time consumed, on average, by each of the previously described operations within a 100*μs* interval when different neuron models are computed, on Preempt-RT and Xenomai 3, respectively.

#### 3.2.3 Validation with hybrid circuits

Proper performance of RTHybrid neuron and synapse models was tested building hybrid circuits as detailed in section 2.2.2. Living neurons from the pyloric CPG were bidirectionally connected through chemical graded synapse models with neuron models. Four trials per model were conducted, each of them five minutes long, with one a minute long control period before and after the hybrid circuit interaction. The first two trials were performed with a sampling frequency of 10kHz, thus cycle interval duration was 100*μs*. The remaining two trials were run at 20kHz, and the interval was 50*μs* long. Any latency value exceeding that limit was considered a real-time failure. All four trials per model were repeated both in Preempt-RT and Xenomai 3, without core isolation.

Connections in these hybrid circuits mimicked graded chemical synapses with fast and slow dynamics (Golowasch et al., 1999). The connection from the model to the living neuron was built with a slow synapse.

**Figure 7.**
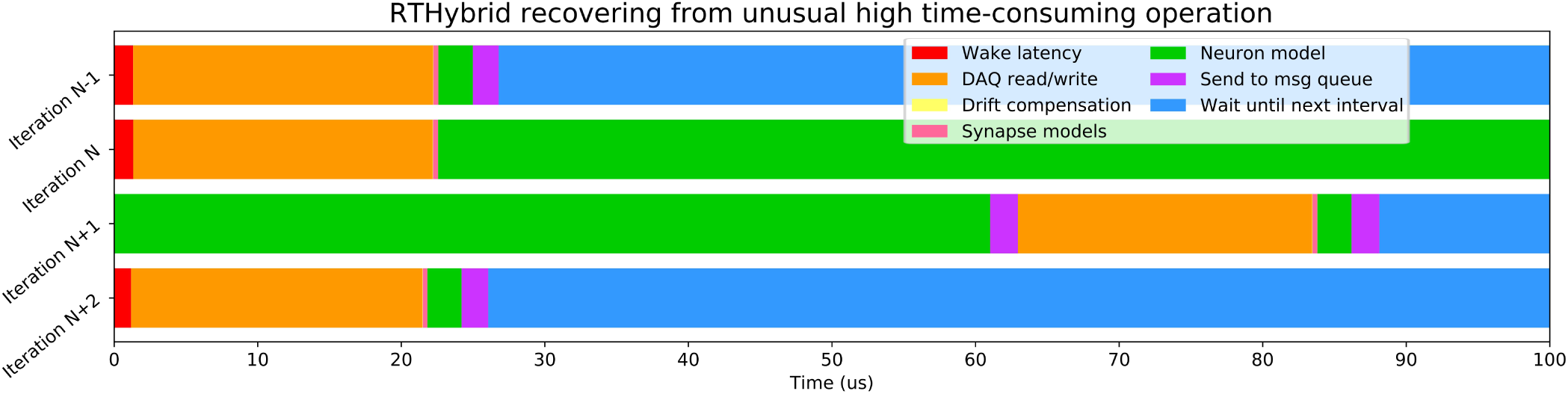
Illustration of how RTHybrid recovers from an unusual high time-consuming operation. In this case, during iteration N of a test on Preempt-RT with Wang model (Wang, 1993), the computation time of the neuron model was abnormally high, reaching almost 140*μs* and, therefore, exceeding the real-time constrains. RTHybrid handles this kind of situation by taking time from iteration *N* + 1 to finish the pending operations from iteration *N*, and reducing or skipping the waiting time, so on iteration *N* + 2 the proper behaviour of the system is restored.

A fast graded synapse was used for the connection from the living neuron to the model. Synapses were inhibitory, resulting in a rhythmic antiphase behaviour between both neurons. The slow synapse had a conductance of *g* = 0.02*μS*, release threshold of *V_th_* = 15% of the maximum amplitude range and kinetic parameters *k*_1_ = 14.0 and *k*_2_ = 4.0. The conductance of the fast synapse was *g* = 0.4*μS*, except for the Hindmarsh-Rose model that had *g* = 0.8*μS*, and the release threshold was set to *V_th_* = 25% of the maximum amplitude range for all but the Ghigliazza-Holmes model neuron, which used *V_th_* = 50%. Input voltage scaling factor was set to 1000 and output current/voltage conversion factor to 1*nA/V*. Neuron model parameters were set to produce bursting behaviour and the duration of each burst was set to automatically match the living neuron activity. Signal amplitude and temporal scaling was automatically performed by RTHybrid calibration algorithms (Reyes-Sanchez et al., 2017).

Figure 8 shows the validation tests results for the Rulkov and Izhikevich models, which are the ones with simpler mathematical descriptions, and Fig. 9 refers to Hindmarsh-Rose and Ghigliazza-Holmes models. For each model there are four panels displaying different information about the hybrid interaction. The worst performance trials for each model, understood as those with higher latency values, were selected for the analysis. Recorded membrane potential from both neurons is displayed in panel A, scaled to the living neuron range, portraying the robust antiphase rhythm achieved due to the bidirectional inhibitory graded chemical synapse used in all cases, as shown at panel B. Panel C represents Preempt-RT test latency values, showing that in all cases the 100*μs* limit established for 10kHz trials, represented by the right-most red vertical line, was fulfilled. However, this RTOS was not able to keep 20kHz constrains, indicated by the left-most red vertical line, during these experiments. This was not the case for Xenomai 3, portrayed at panel D, whose latency values were far under the 50*μs* barrier set during 20kHz trials. Due to this outcome, the results displayed correspond to Preempt-RT 10kHz and Xenomai 20kHz trials.

**Figure 8.**
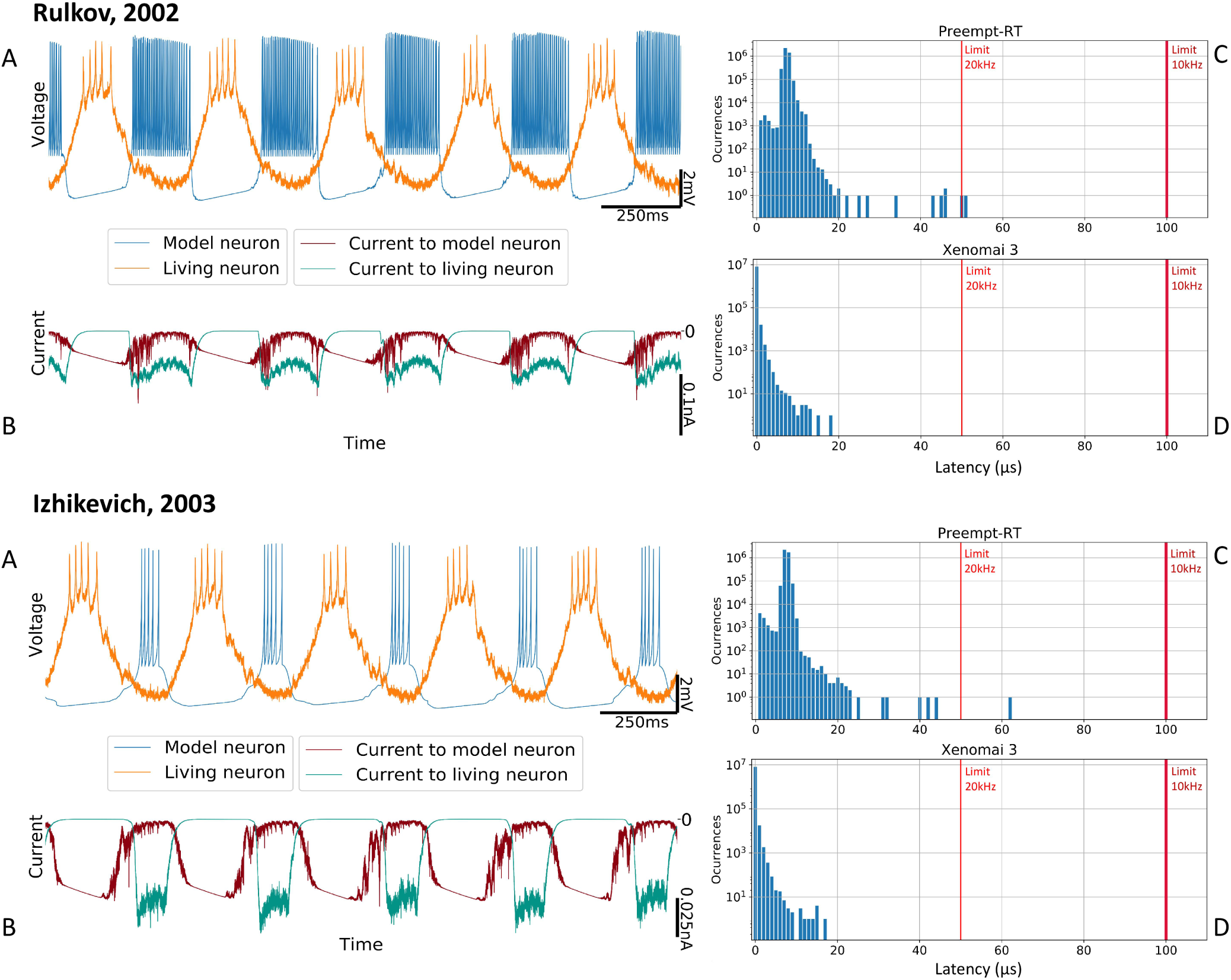
Hybrid circuit built with a pyloric CPG neuron and a simulated neuron using models from (Rulkov, 2002) and (Izhikevich, 2003). For both models, each of the four panels represent: A) membrane potential of the neurons (adapted to the living neuron range) B) synaptic currents C) Preempt-RT latency values D) Xenomai 3 latency values. On the latency figures, left-most red line represents the 20kHz limit and the right-most, the 10kHz limit.

**Figure 9.**
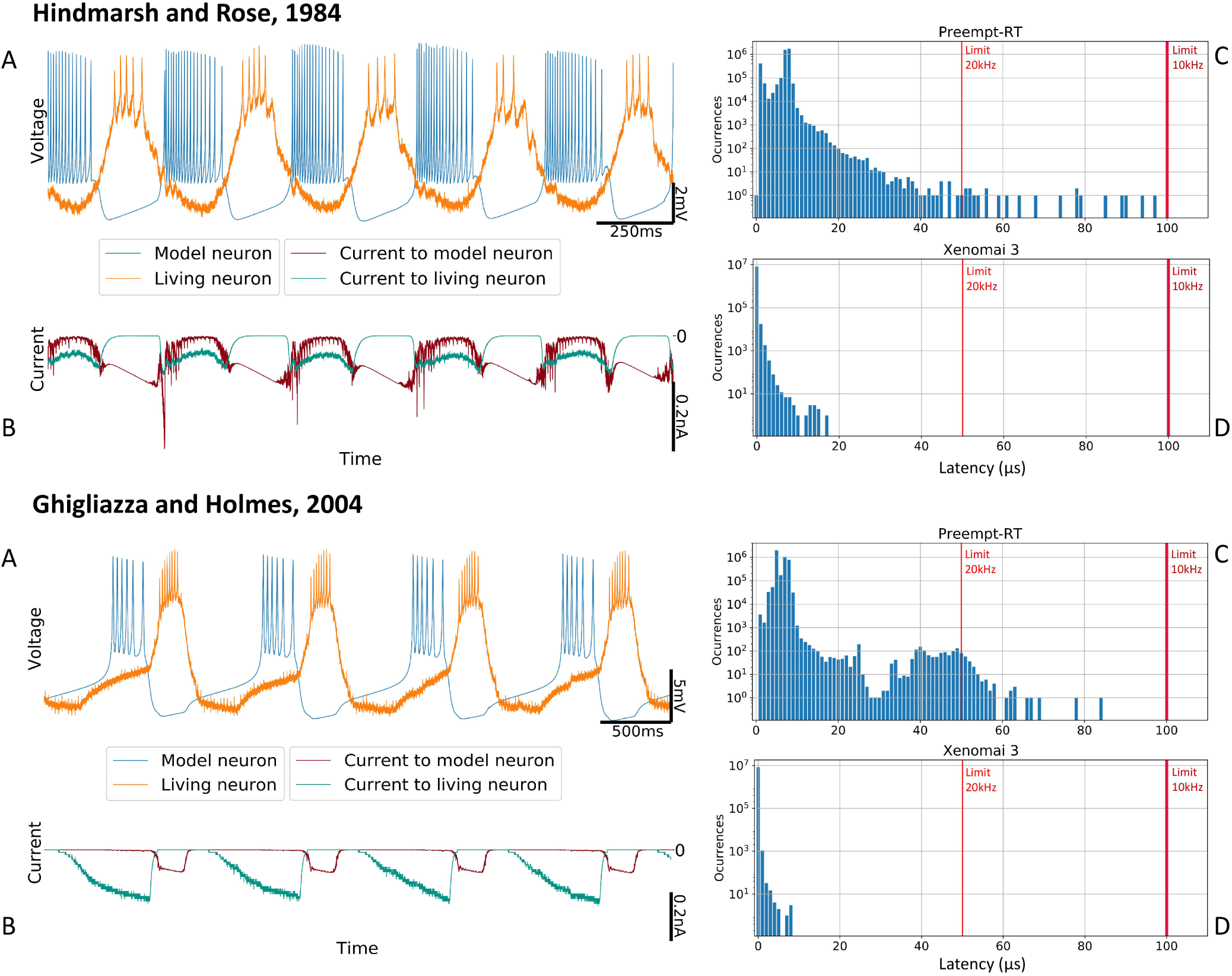
Hybrid circuit built with a pyloric CPG neuron and a simulated neuron using models from (Hindmarsh and Rose, 1984) and (Ghigliazza and Holmes, 2004). For both models, each of the four panels represent: A) membrane potential of the neurons (adapted to the living neuron range) B) synaptic currents C) Preempt-RT latency values D) Xenomai 3 latency values. On the latency figures, left-most red line represents the 20kHz limit and the right-most, the 10kHz limit.

## 4 DISCUSSION

Characterization and control of neural systems dynamics, as well as experimental protocol automation can largely benefit from the use of closed-loop techniques, and specifically from hybrid circuits built by connecting model neurons and synapses to living cells. RTHybrid provides a software neuron and synapse model library aimed to build hybrid circuits in an easy and user-friendly manner. Developed for Linux, this program is open-source and can be downloaded for free from www.github.com/GNB-UAM/RTHybrid, where installation and user manuals can also be found. RTHybrid has a simple GUI to design and configure the experiments. This tool also incorporates a command-line mode where configuration XML files can be loaded, a useful feature for experiment automation and scripting.

Temporal precision requirements for closed-loop interactions in RTHybrid is fulfilled by using hard real-time software technology. An extensive analysis of available RTOS was conducted to select the most suitable platforms to implement RTHybrid. The software has been developed to run over Preempt-RT and Xenomai 3 frameworks for Linux due to their balance between performance and accessibility. RTHybrid model library includes a wide variety of neuron models: from computationally-unexpensive paradigms, such as (Rulkov, 2002; Izhikevich, 2003), to realistic conductance-based Hodgkin-Huxley type (Ghigliazza and Holmes, 2004; Wang, 1993). The library also contains several synapse models: from simple gap junctions implementations to configurable graded chemical synapses (Golowasch et al., 1999). All of them are adapted to work under hard real-time restrictions. Moreover, calibration algorithms are integrated in the library to automate the adaptation of model amplitude and time scales to the living neuron behavioural range (Reyes-Sanchez et al., 2017).

Models currently included in the RTHybrid library are suitable for a wide variety of hybrid circuit experiments implemented using dynamic clamp. Beyond electrophysiological protocols, RTHybrid can also be easily generalized to drive open- and closed-loop interactions in optogenetic and drug microinjection paradigms. Future development in parallelization and GPU computing will be considered to implement large scale network or highly-realistic biophysical models.

Many researchers and laboratories overlook closed-loop techniques despite their advantages due to the difficulties in the installation and use of the required technology. With RTHybrid, we aim to encourage the use of open-source, standardized and user-friendly real-time software tools, available in different platforms, to facilitate the implementation of closed-loop experimentation in neuroscience research.

## Footnotes

1 Source code of the latency test: www.github.com/RoyVII/Latency_tests.git

2 stress software website: www.people.seas.harvard.edu/~apw/stress/

3 Ubuntu 16.04 with Xenomai 3.0.5 Live CD/USB: www.mega.nz/#!CPwCRIJQ!jWVGb08wjSP-02-kiv7KK_bKzcaPERWBp7MjH6coXVs

4 Source code for RTHybrid: www.github.com/GNB-UAM/RTHybrid

5 RTHybrid User Manual: www.github.com/GNB-UAM/RTHybrid/blob/master/docs/RTHybrid_User_Manual.pdf

6 Qt5 website: www.qt.io/

7 libxml2 library website: http://xmlsoft.org/

